# Serum proteins can successfully predict self-reported ethnicity: Implications for precision-medicine

**DOI:** 10.1101/221614

**Authors:** Ram J Bishnoi, Raymond Palmer, Donald R Royall

## Abstract

The effects of inter-individual variability on disease treatment and prevention are important to the goals of “precision medicine” ^1^. In biomedical research, consideration of racial or ethnic differences allows generation and exploration of hypotheses about interactions among genetic and environmental factors responsible for differential medical outcomes. The US National Institutes of Health, therefore recommends adequate participation of subjects from ethnic minority groups in research studies. Nevertheless, considerable debate has focused on validity of race or ethnicity as biological construct ^2^. Inconsistent definition of race/ethnicity and insignificant genetic variations between ethnic groups have invited disregard to this construct ^3^. On the contrary, differences in prevalence, expression and outcomes of various diseases among ethnic groups argue for continued and focused attention to ethnicity as important predicting variable. In context of Alzheimer’s disease (AD), we have previously reported that ethnicity does moderates the proteomic markers of dementia ^4^. Here, we attempted to classify and predict selfreported ethnicity (Hispanic or non-Hispanic white, [NHW]) using a limited serum profile of 107 proteins. Random Forest (RF) classification method was able to discriminate those two ethnicities with 95% accuracy and could successfully predict ethnicity in an independent test-set (Area under ROC curve: 0.97). Variable selection method led to a condensed set of six proteins which yielded comparable classification and prediction accuracy. Our results provide preliminary evidence for proteomic variability between ethnic groups, and biological validity of ethnicity construct. Moreover, they also offer an opportunity to exploit these differences towards the objectives of precision medicine.

## Main

A secondary objective of the Texas Alzheimer’s Research & Care Consortium (TARCC) project, an ongoing longitudinal cohort study aimed at examining the role of clinical and biological markers of AD, is to investigate ethnic variability with regard to clinical features and biomarkers. Majority of participants in TARCC belong to Hispanics (Mexican-Americans; MA) or non-Hispanic whites (NHW), two major ethnic groups of Texas. Like most research studies, ethnicity in TARCC was self-reported. Five hundred and thirty cognitively healthy elderly individuals (312 Hispanics and 218 NHWs) who were recruited as control subjects in TARCC study were used for this analysis. Two ethnic groups differed significantly in terms of age, education, prevalence of diabetes and obesity, and an “omnibus” dementia severity metric (i.e., “δ”) ^4^ (Table 1). Although nominally non-demented, these control subjects were recruited for a convenient sample study and two groups differed with regard to “δ“. After removing the variance related to available confounding variables, 107 serum proteins from baseline visit and RF algorithm classified two ethnicity classes in a training set (comprising 75% of total the sample) with 95% accuracy. In the test-set (the remaining 25% of total sample), this classification model provided good predictive accuracy with specificity and sensitivity of 93.10% and 97.33% respectively with area under Receiver Operating Characteristic (AUC-ROC) curve of 0.97. Twenty most discriminatory variables, identified by variable importance (VI) measure could predict ethnicity with AUC-ROC of 0.99, sensitivity of 97.33% and specificity of 93.10%. . Finally, in order to limit the number of classifying variables to facilitate future replicability, a variable selection method provided a set of six protein variables, which provided a comparable predictive accuracy with ROC AUC of 0.99, sensitivity of 94.87% and specificity of 94.55%. Results from all three models are summarized in Table 2.

**Table 1.** Demographic and clinical characteristics of participants

|  | Hispanic | NHW | P value |
| --- | --- | --- | --- |
| N | 312 | 218 |  |
| Age (mean, SD) | 64.99, 7.94 | 70.43, 8.56 | <0.0001 |
| Gender(% female) | 62.5 | 67 | 0.3342 |
| Education (mean, SD) | 11.27, 4.66 | 15.60, 2.62 | <0.0001 |
| APOE e4(%) | 19.2 | 26.2 | 0.0747 |
| Hypertension(%) | 60.9 | 57.8 | 0.5316 |
| Diabetes(%) | 33 | 11.5 | <0.0001 |
| Obesity(%) | 49.4 | 22.9 | <0.0001 |
| Hyperlipidemia (%) | 55.4 | 49.1 | 0.1628 |
| Dementia severity metric, $\delta$ (Mean, SD) | -0.40, 0.38 | -0.72, 0.37 | <0.0001 |

**Table 2.**
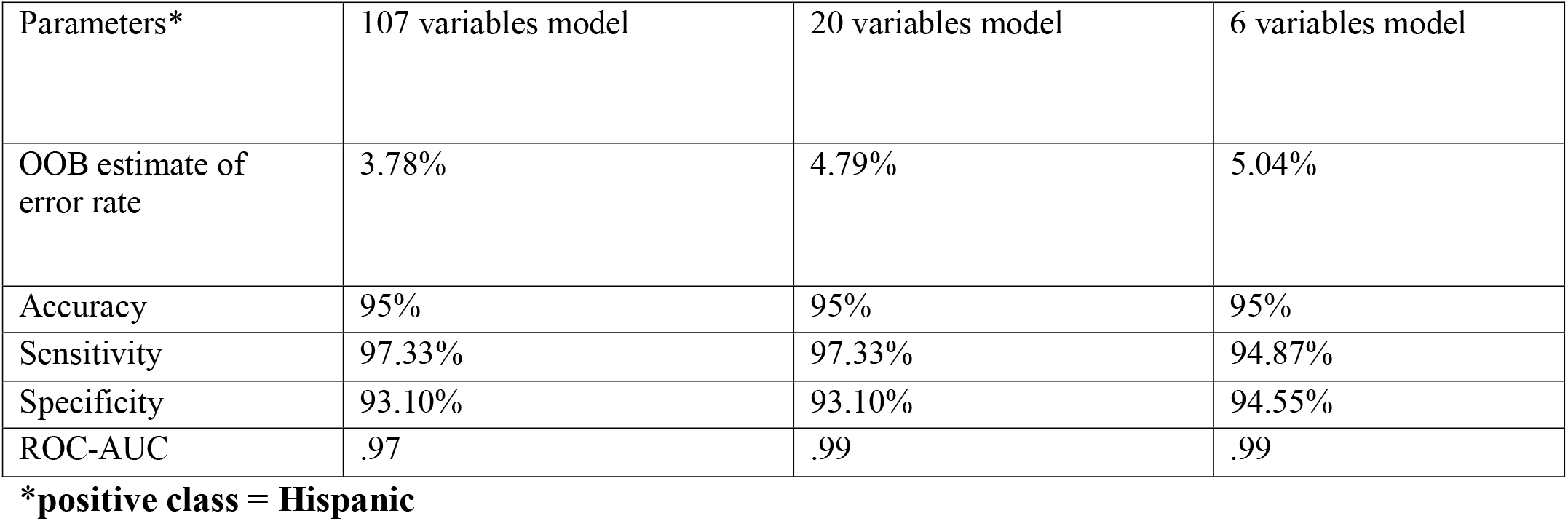
Classification and prediction results. Out of bag (OOB) error rate estimated prediction error of random forest models utilizing bootstrap aggregating to training set. Classification performance of a random forest model on test set was assessed by classification matrix and area under the receiver-operating characteristic curve (AUC). The performance was explored for three different sets of explanatory variables.

These results support the potential validity and utility of ethnicity as a biological construct ^5^. Even though Hispanic group is a heterogeneous mixture of various ancestries ^6^ and not as mutually exclusive from NHWs as for example NHWs and east-Asians, serum protein’s ability to distinguish it from NHWs highlight importance of ethnicity in not only biomarker research but also disease pathophysiology. Previously, ethnicity related differences in expression of several blood-based biomarkers, including cytokines ^7–9^ have been reported from TARCC ^10–13^, Multi-Ethnic Study of Atherosclerosis (MESA) ^14–21^ and National Health and Nutrition Examination Survey (NHANES) ^22–29^. Combined with our results, these findings lay a common theme that supports our hypothesis that ethnicity does impact the circulating levels of proteins. The mechanism(s) that mediate those differences remain unclear. In the simplest application, the biomarkers of ethnicity can be viewed as biological parameters that are themselves significantly correlated with an ethnicity and not necessarily responsible for ethnic groupings. Therefore, the resulting protein profile may not be the biological determinants of ethnicity per se but a reflection instead of cross-ethnic differences in lifestyle or disease processes. These differences among ethnic groups may be responsible for disparate disease-susceptibility, treatment response and/or outcomes. Our results strongly emphasize the importance of self-reported ethnicity information in research and underlines the need for validation of research results across ethnicities.

Multiple prior attempts to measure the correspondence between self-reported ethnicity class and genetic clusters have provided mixed results ^2,3,30,31^. Human genome sequencing project and related studies concluded that ethnicity is a weak surrogate of various genetic and non-genetic factors in correlations with health status ^32^. In fact, individuals from different ethnicity were found to be genetically similar than individuals from their own ethnicity ^3,33^. Surprisingly, a limited proteomic assay which was not pre-selected to predict ethnicity or to develop ethnicity specific traits, could precisely differentiate & predict ethnicity. Complete human proteome may provide greater precision and possibly provide ethnicity specific proteomic compositions. Our results therefore emphasize importance of proteomic research and endorse the hypothesis that proteomics is not just the new genomics ^34^ and include phenotypic information from protein posttranslational modifications, protein interactions, metabolite abundance, and, protein stoichiometry ^35^.

Classification models using a reduced set of the six most important differentiating proteins resulted in comparable predictive accuracy. These six proteins were-Connective tissue growth factor (CTGF), interferon (IFN) gamma, interleukin (IL)-12p40, IL-13, transforming growth factor (TGF) alpha, and thrombospondin 1 (THBS1). Relative expression of each of these six proteins is shown in fig 3. IL13 levels were relatively higher in Hispanic group, although not statistically significant (p=0.12). All other proteins were over-expressed in NHW group, levels were statistically significant with exception of IL12p40 (p=0.83). These reflect mean differences, however, Figure 3 also suggests clear cross-group differences in these proteins’ distributions. Protein-protein interactions (PPI) network of identified proteins indicated that these selected proteins were enriched for various processes including regulation of protein phosphorylation, cell activation and intracellular signal transduction (Table 3). CTGF and TGF-alpha are epidermal growth factor receptor (EGFR; a member of tyrosine kinase family) ligands which controls cellular functions during development and homeostasis through protein phosphorylation ^36^. EGFRs have been found to be dysregulated in several epithelial cancers like lung, colon etc and known to vary among ethnic groups ^37,38^. Among these epithelial cancers, ethnicity influence response to treatment and mortality. For example, response to EGFR tyrosine kinase inhibitors (e.g. geftinib) treatments in lung cancer can be predicted by ECFR ligands levels in blood and ethnicity ^39^. Among ethnicities, Hispanic and Asian groups respond better to these treatments compared to NHWs ^39^. Lower levels of EGFR ligands like CTGF & TGF-alpha may play a role in comparatively better response to EGFR inhibitors in Hispanics. Our results do not provide any direct evidence to this hypothesis but support the arguments for further investigations into this direction. If serum levels of ligands like TGF alpha can predict response to treatments, we can anticipate rapid development in precision medicine based on serum proteomic assays. Also, if EGFR activity is proven to be significantly different among ethnic groups, ethnicity can provide individual variability to patients which is key to precision medicine.

**Fig 1.**
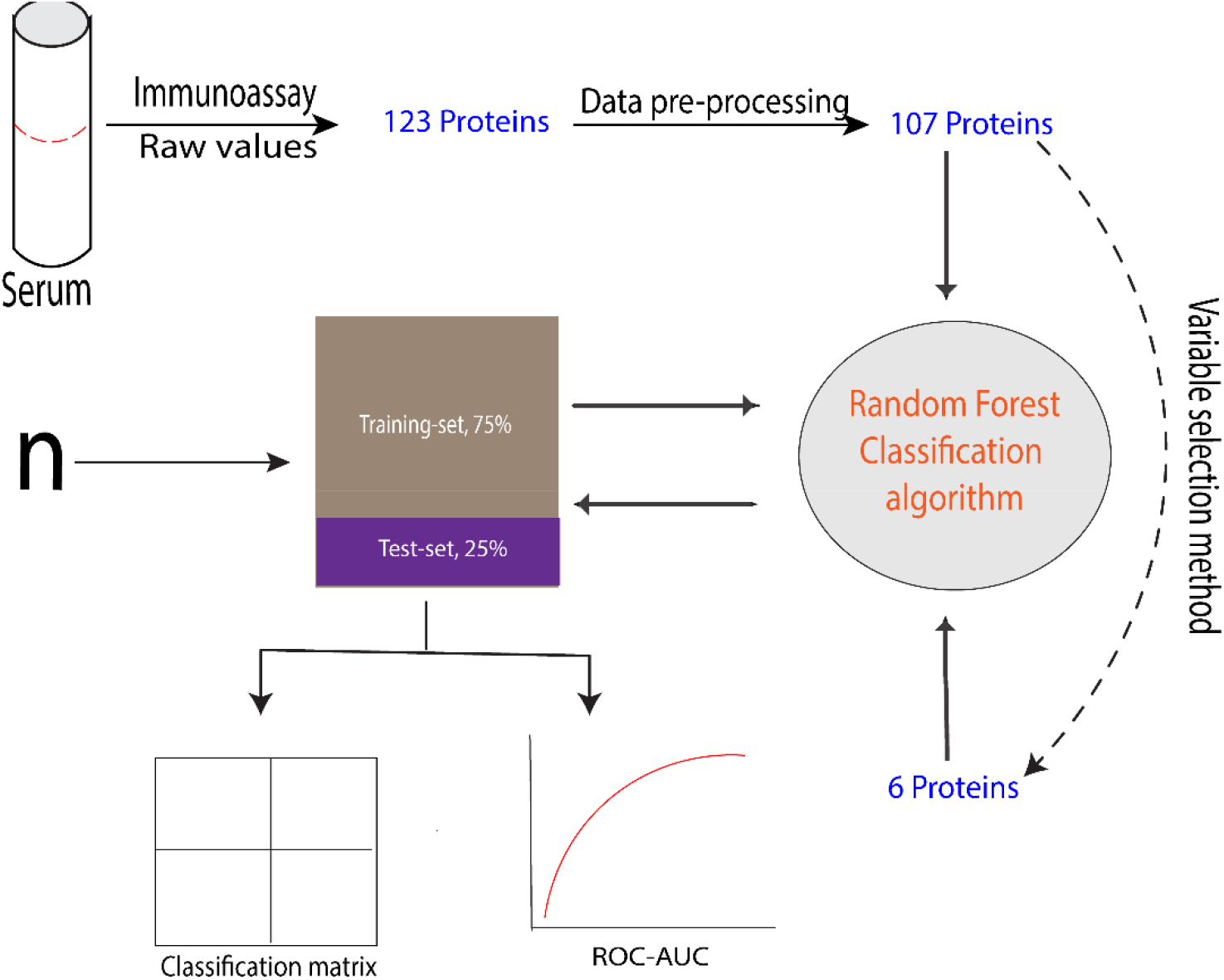
Overview of methodology

**Fig 2.**
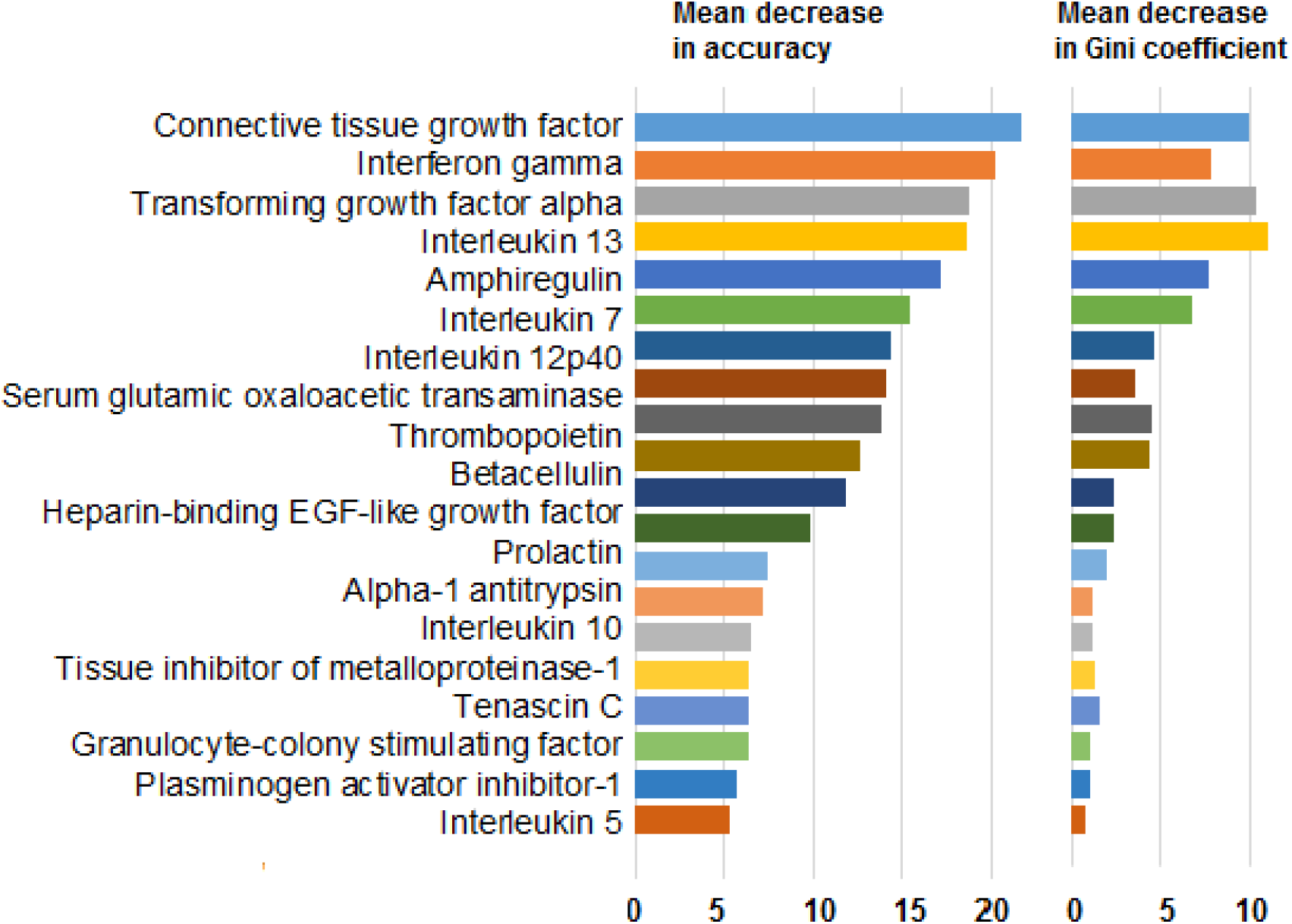
Variable importance for Random Forest classification model, ranked based on mean decrease in accuracy. Only top 20 variables are shown here along with their relative Gini coefficients.

**Fig 3.**
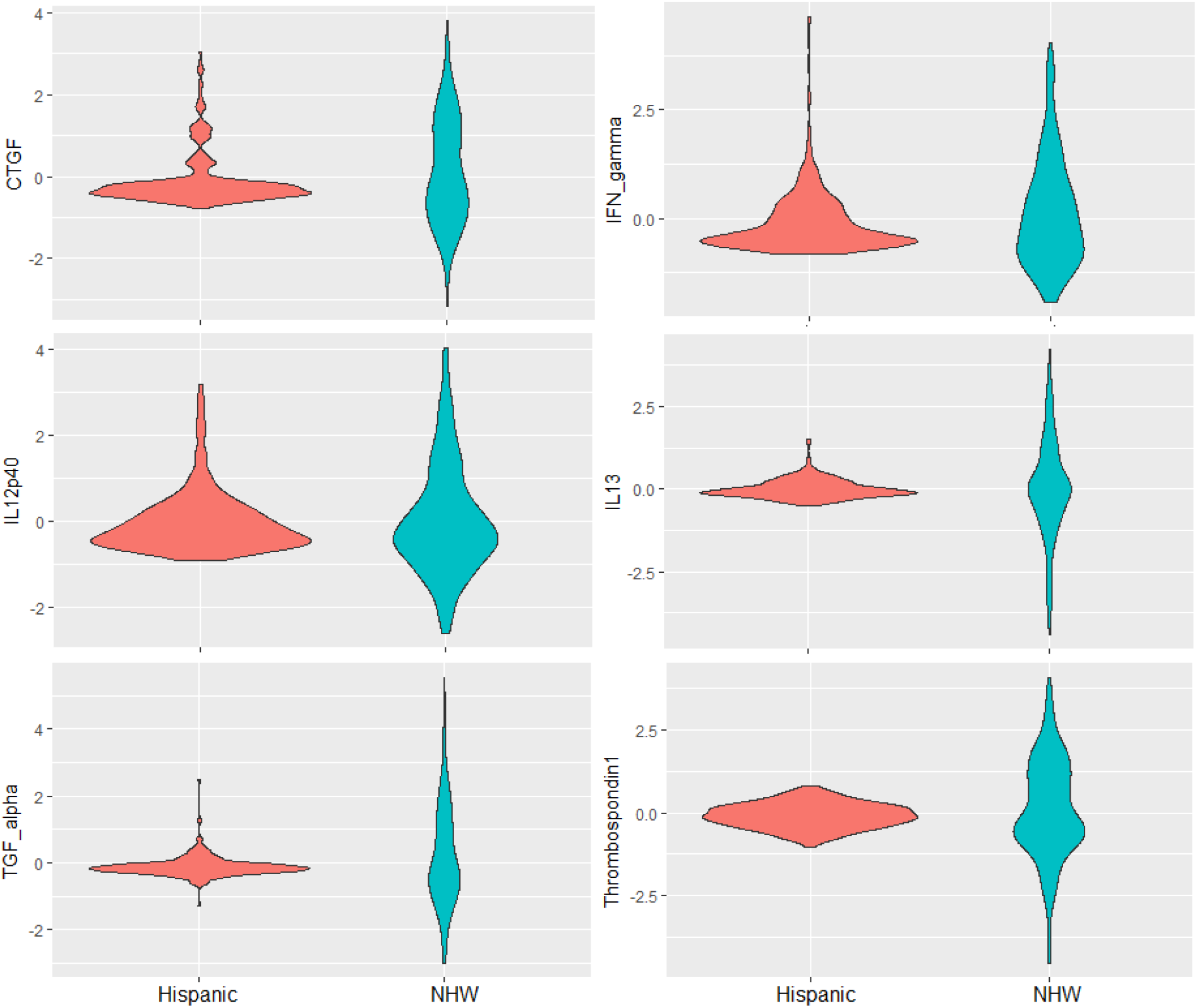
Differential expression of most significant proteins differentiating Hispanic and NHW groups in Random Forest classification model is shown in violin plot. Mean level of IL13 was higher in Hispanic group but difference was not significant (p=.97) while all other proteins were overexpressed in NHW group. CTGF (p=.04), IFN gamma (p=.03), TGF alpha (p=.00) and Thrombospondin 1 (p=.02) were significantly different in two groups. Difference in mean level of IL12p40 (p=.76) was insignificant.

**Fig 4.**
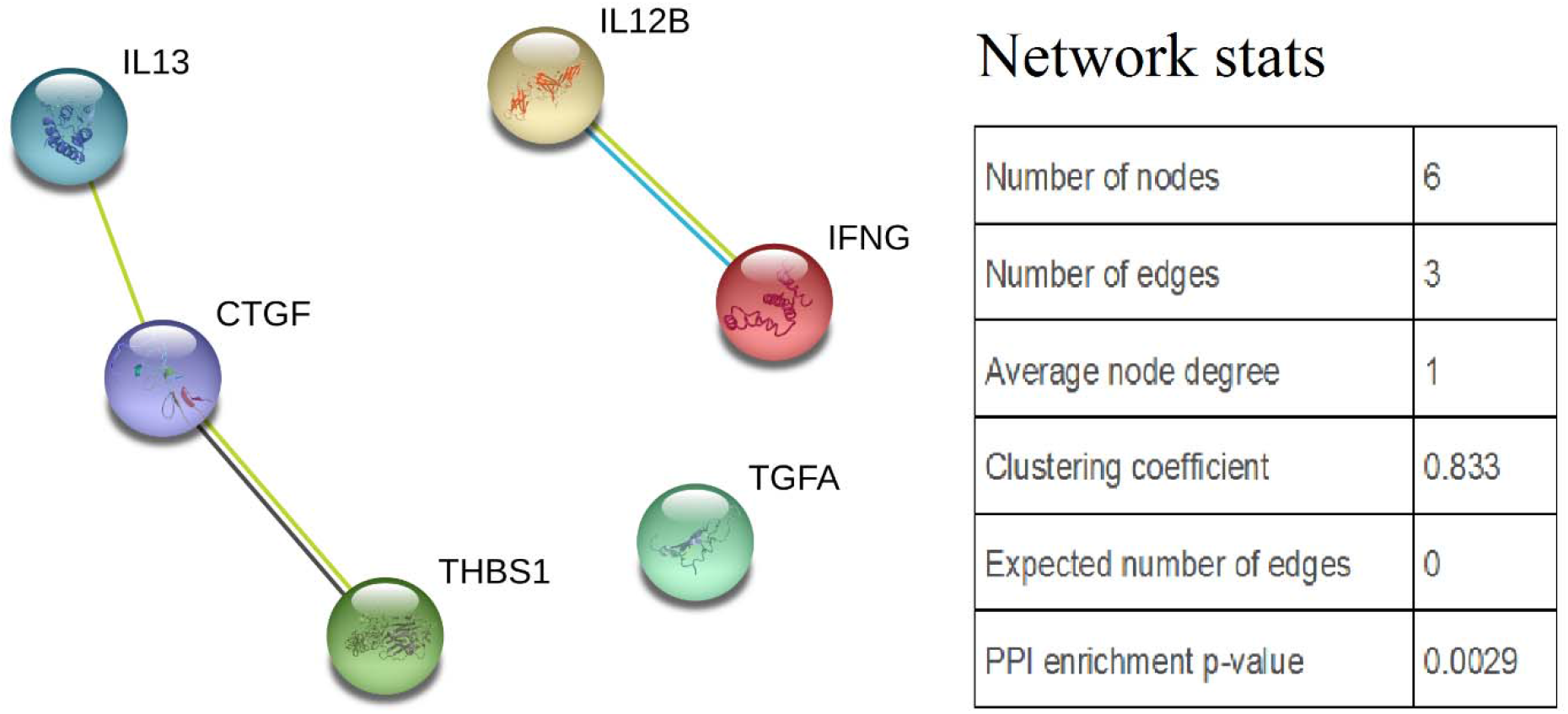
Protein-protein interaction networks of top six classifiers of two ethnicities (Hispanics and NHWs).

**Table 3.** Functional enrichments in protein-protein interactions network

| Pathway description | Count in gene set | False discovery rate |
| --- | --- | --- |
| **Biological processes (GO)** |  |  |
| positive regulation of protein phosphorylation | 6 | 9.93e-06 |
| positive regulation of cell activation | 5 | 9.93e-06 |
| positive regulation of intracellular signal transduction | 6 | 9.93e-06 |
| positive regulation of peptidyl-tyrosine phosphorylation | 4 | 4.61e-05 |
| regulation of apoptotic process | 6 | 4.68e-05 |
| **Molecular functions (GO)** |  |  |
| Receptor binding | 6 | 6.93e-05 |
| Cytokine receptor binding | 3 | 0.0399 |

All six selected proteins have been previously reported to associate with dementia phenotype “δ” in ethnicity-adjusted models from TARCC cohort ^41,42^. Details about concept and utility of δ is can be obtained from past publications ^43–45 46^. There are cross-ethnic differences in the δ-scores of non-demented TARCC participants, and in δ’s serum protein biomarkers ^12^. Although the reported results were adjusted for δ, the appearance of δ-specific biomarkers among the most discriminating set of ethnicity related biomarkers may suggest cross-ethnic differences in general intelligence.

Results of this study are also significant in context of “Hispanic paradox”, an epidemiological paradox observed in southwestern states of US including Texas ^40^. Socioeconomically, Hispanics were disadvantaged and resembled African-Americans but their health status was comparable to NHWs. Precise biological basis to this phenomenon is unclear and several theories have been proposed to explain this paradox i.e. Salmon bias effect, selective migration, better social support etc. Differential response to EGFR inhibitors in lung cancer and expression of EGFR ligands in Hispanics is probably an example of biological factors which offset the impact of socio-economic adversities. Our study provides merely a direction for further research focused on ethnicity specific biological differences.

Our analysis has notable limitations. Even though we included a relatively large sample size and more than 100 serum proteins, quality control measures to remove batch effects, and adjustment for demographic and clinical factors, these results require validation in another independent sample. A complete assay of serum proteomics could be more valuable in such analysis. Also, all proteins were measured cross-sectionally and it was not possible to verify if ethnicity specific profile were stable over time. Finally, patient population used in this study included elderly individuals and these results need validation in younger adults ^47^.

In summary, we provide proof of concept for the accurate prediction of self-reported ethnicity from a set of serum proteins. There exists significant support in literature for differential protein expression by ethnic group. These may inform disparities in disease prevalence, severity or treatment response and may be of value to the development of “precision medicine”.

## Methods

Full design of study is outlined in supplementary fig 1. Data were derived from Texas Alzheimer’s Research & Care Consortium (TARCC) project, an ongoing longitudinal cohort study aimed at examining the role of clinical and biological markers in the development and progression of Alzheimer’s disease (AD). Each participant in this study provided written informed consent prior to enrollment, evaluation and biomarker blood draw. Detailed description of the TARCC study design can be found elsewhere ^48,49^.

The demographic and clinical characteristics of patients are summarized in supplementary Table 1. For this analysis, 530 cognitively healthy elderly individuals (Hispanics n= 312, NHWs n= 218) who participated as control subjects in TARCC study were considered. Each participant underwent a standardized evaluation at respective TARCC sites on baseline and then annually. At baseline, all subjects underwent clinical interview, neuopsychological assessment and biomarker blood draw. A consensus group for the respective site determined their cognitive status as cognitively healthy individuals. Study participants with serious medical illness were not considered for the study. Inclusion and exclusion criteria have been discussed previously in details ^48^.

## Proteomic profiling

Non-fasting samples were collected with 10mL serum-separating (tiger-top) vacutainers tubes at the baseline visit. Samples were allowed to clot at room temperature for 30 minutes in a vertical position before being centrifuged at 1300 × g for 10 minutes. Next, 1mL aliquots were transferred into polypropylene cryovial tubes labeled with freezerworks barcodes and placed into −80° C freezers for storage until use.

Serum samples were shipped to Myriad RBM, a CLIA-certified biomarker testing Laboratory based in Austin, TX, USA. The samples were assayed on the luminex-based HumanMAP 1.0 platform. Over 100 proteins were quantified utilizing fluorescent microspheres with protein-specific antibodies. Information regarding the least detectable dose (LDD), inter-run coefficient of variation, dynamic range, overall spiked standard recovery, and cross-reactivity with other HumanMAP analytes can be obtained online (https://rbm.myriad.com/products-services/humanmap-services/humanmap/).

## Statistical analysis

### Data pre-treatment

Analytes with >15% missing data or value below the lower detection limit (LDL), were excluded from analysis. Out of 124, 107 analytes were included for further analysis and missing values were imputed with k-nearest neighbor (KNN) method with 10 neighbors per analyte based on Euclidean distance. KNN is an imputation method that accounts for the local similarity of the data by identifying similar analytes with similar peak intensity profiles via a distance ^50^. Next, data were log(10)-transformed and normalized to reduce systemic variance among the data and improve the performance of downstream analysis. Correlation and colinearity was tested to ensure the variables are independent. Pearson correlation was high for some variables like different interleukins which is not completely unexpected, however the Variance Inflation Factors (VIF) were less than 10 for all variables which is accepted threshold for multi-collinearity.

Following potential confounders were considered: age, sex, study site, batch number, APOE status, educational achievement and history of hypertension, diabetes, hyperlipidemia and obesity. Each analyte variable was regressed onto covariates to adjust for the confounding effects on analytes. The standardized residuals (mean of zero and standard deviation of one) from the linear regression were then used in further analysis to classify and predict ethnicity.

### Classification methods

Dataset was divided into a training set (75%) and an independent test dataset (25%), and we trained a Random Forest (RF) in the training set of Hispanic and NHW subjects using their proteomic profiles. We evaluated its performance using cross-validation on test-set and scored predictive power in a receiver operating characteristic (ROC) analysis.

We selected Random Forest (RF) model ^51^, an increasingly popular method in machine learning that characterizes structure in high dimensional data while making no distributional assumptions about the response variable or predictors. RF model (using *randomForest, an R* package) uses an ensemble of classification trees was employed to develop a Hispanic versus NHW classifier from the training set. RF builds a forest with user defined number of regression trees (ntree) and number of predictors sampled for splitting at each node (mtry); we used 500 classification trees and mtry=Vn. Trees are built by recursive binary partitioning of observations into subsets and each tree is asymptomatically unbiased. A portion of the observations is excluded from the tree building process (i.e. out-of-bag or OOB data) for each individual tree to be used as an independent estimate of the prediction accuracy. At each node in a tree, the variables used to develop that tree are tested for the split in the data that achieves greatest reduction in error. RF used this ensemble of trees to make predictions of class based on majority votes to a given sets of parameters. During this process, “variable importance (VI)” is assessed that measures the relative predictive influence of individual variables on the classification model. VI based on permutation accuracy importance measure is most advanced measure because of its ability to evaluate the variable importance by the mean decrease in accuracy using the internal out-of-bag (OOB) estimates while the forests are constructed ^51–53^.

### Variable selection

Although RF provides a measure of variable importance, it does not automatically choose the optimal number of variables that can yield the good classification accuracy with low misclassification error rate. To identify a set of limited number of variables, we employed two strategies. First, we used the top-20 variables based on VI measure and second, we employed a backward elimination of variables using OOB errors ^54^ This was performed using R package varSelRF and variable elimination from RF was carried out by successively eliminating the least important variables (with importance as returned from random forest) and minimizing OOB error at the same time. At each iteration, a fraction of variables (default value = 0.2) were excluded from those in the previous forest that lowered computational cost and is coherent with the idea of an “aggressive variable selection” approach. In this process, model resolution gets better as the number of variables considered becomes smaller. After fitting all forests, the selected set of features is the one whose OOB error rate is within u = 1 standard error of the minimum error rate of all forests. This is in agreement with the “1 s.e. rule” commonly used in classification trees literature ^55^. The objective of these strategies was to select optimal subset of variables while keeping an error rate close to that from the original model. These subsets of variables were then used to classify ethnicity groups in training data and tested with independent test set.

### Classification performance

At each step, the performance of the RF was assessed by OOB estimate of error rate ^52,56^ and a separate accuracy assessment on an independent data set. Confusion matrix was subsequently constructed to compare true class with the class assigned by the classifier and to calculate overall accuracy. Area under the curve (AUC) of the receiver operating characteristic (ROC) curve estimated the predictive performance of models ^57,58^.

## Functional analysis of protein-set

Most significant proteins variables derived from variable selection method were assessed for their functional correlates with the STRING (Search Tool for the Retrieval of Interacting Genes/Proteins) database (http://string-db.org) ^59^. STRING allows exploration of how these proteins are inter-related to form protein-protein interaction networks by applying all active interaction sources (experiments, databases and text mining). It can also provide biological functions and specific cellular pathways involving those proteins.

## Acknowledgemnts

### Author contributions

RJB designed experiments, analyzed data and wrote the paper. RMP pre-processed data and assisted data analysis. All authors edited the paper and DRR supervised the research.

This study was made possible by the Julia and Van Buren Parr endowment for dementia-related research and the Texas Alzheimer’s Research and Care Consortium (TARCC) funded by the state of Texas through the Texas Council on Alzheimer’s Disease and Related Disorders. The results have been presented as a poster at the University of Texas Health Science Center, San Antonio Research Day, 2016.

Investigators from the Texas Alzheimer’s Research and Care Consortium: Baylor College of Medicine: Valory Pavlik, PhD, Paul Massman PhD, Eveleen Darby MA/MS, Monica Rodriguear MA, Aisha Khaleeq MD; Texas Tech University Health Sciences Center: John C. DeToledo, MD, Henrick Wilms MD, PhD, Kim Johnson PhD, Victoria Perez, Michelle Hernandez; University of North Texas Health Science Center: Thomas Fairchild PhD, Janice Knebl DO, Sid E. O’Bryant PhD, James R. Hall PhD, Leigh Johnson PhD, Robert C. Barber PhD, Douglas Mains DrPH, Lisa Alvarez, Adriana Gamboa; University of Texas at Austin: John Bertelson MD; University of Texas Southwestern Medical Center: Perrie Adams PhD, Munro Cullum PhD, Roger Rosenberg MD, Benjamin Williams MD, PhD, Mary Quiceno MD, Joan Reisch PhD, Linda S. Hynan PhD, Ryan Huebinger PhD, Janet Smith BS, Barb Davis MA, Trung Nguyen MD, PhD; University of Texas Health Science Center – San Antonio: Donald Royall MD, Raymond Palmer PhD, Marsha Polk; Texas A&M University Health Science Center: Farida Sohrabji PhD, Steve Balsis PhD, Rajesh Miranda, PhD; University of North Carolina: Kirk C. Wilhelmsen MD, PhD, Jeffrey L. Tilson PhD, Scott Chasse, PhD.

### Competing financial interests

Drs. Royall and Palmer have disclosed the results of these analyses to the University of Texas Health Science Center at San Antonio (UTHSCSA), which has filed patent application 2012.039.US1.HSCS and provisional patents 61/603,226 and 61/671,858 relating to the latent variable δ’s construction and biomarkers.

Dr. Royall reports research funding from the Department of Defense (DoD), Eisai and Novartis. Drs. Royall and Palmer receive additional funding from the NIH. Dr. Bishnoi reports no biomedical financial interests or potential conflicts of interest.

### Materials and correspondence

Correspondence should be addressed to RB. Data are from the Texas Alzheimer’s Research and Care Consortium (TARCC) study. Requests for TARCC data may be sent to www.txalzresearch.org or to the below contact: Robert Barber, PhD, Associate Professor, Executive Director, Institute for Molecular Medicine, UNT Health Science Center.

## References

1 Council, N. R. Toward precision medicine: building a knowledge network for biomedical research and a new taxonomy of disease. (National Academies Press, 2011).

2 Burchard, E. G., Ziv, E., Pérez-Stable, E. J. & Sheppard, D. The importance of race and ethnic background in biomedical research and clinical practice. The New England journal of medicine 348, 1170 (2003).

3 Risch, N., Burchard, E., Ziv, E. & Tang, H. Categorization of humans in biomedical research: genes, race and disease. Genome biology 3, comment2007. 2001 (2002).

4 Royall, D. R., Palmer, R. F. & Texas Alzheimer’s Res Care, C. Ethnicity Moderates Dementia’s Biomarkers. Journal of Alzheimers Disease 43, 275–287, doi:10.3233/jad-140264 (2015).

5 Bamshad, M., Wooding, S., Salisbury, B. A. & Stephens, J. C. Deconstructing the relationship between genetics and race. Nat Rev Genet 5, 598–609, doi:10.1038/nrg1401 (2004).

6 Stephens, J. C. et al. Haplotype variation and linkage disequilibrium in 313 human genes. Science 293, 489-493, doi:10.1126/science.1059431 (2001).

7 Carroll, J. F. et al. Impact of Race/Ethnicity on the Relationship Between Visceral Fat and Inflammatory Biomarkers. Obesity 17, 1420–1427, doi:10.1038/oby.2008.657 (2009).

8 Stowe, R. P., Peek, M. K., Cutchin, M. P. & Goodwin, J. S. Plasma cytokine levels in a population-based study: relation to age and ethnicity. J Gerontol A Biol SciMed Sci 65, 429–433, doi:10.1093/gerona/glp198 (2010).

9 Majka, D. S. et al. Physical activity and high-sensitivity C-reactive protein: the multi-ethnic study of atherosclerosis. Am J Prev Med 36, 56–62, doi:10.1016/j.amepre.2008.09.031 (2009).

10 O’Bryant, S. E. et al. Biomarkers of Alzheimer’s Disease Among Mexican Americans. Journal of Alzheimers Disease 34, 841–849, doi:10.3233/jad-122074 (2013).

11 Royall, D. R., Palmer, R. F. & Consortium, T. A. s. R. a. C. Does ethnicity moderate dementia’s biomarkers? Neurobiol Aging 35, 336–344, doi:10.1016/j.neurobiolaging.2013.08.006 (2014).

12 Royall, D. R. & Palmer, R. F. Ethnicity moderates dementia’s biomarkers. J Alzheimers Dis 43, 275–287, doi:10.3233/JAD-140264 (2015).

13 Royall, D. R. & Palmer, R. F. Thrombopoietin is associated with δ’s intercept, and only in Non-Hispanic Whites. Alzheimers Dement (Amst) 3, 35–42, doi:10.1016/j.dadm.2016.02.003 (2016).

14 Bradley, R. D. et al. Associations between gamma-glutamyltransferase (GGT) and biomarkers of atherosclerosis: The multi-ethnic study of atherosclerosis (MESA). Atherosclerosis 233, 387–393, doi:10.1016/j.atherosclerosis.2014.01.010 (2014).

15 Jenny, N. S. et al. Associations of inflammatory markers with coronary artery calcification: Results from the Multi-Ethnic Study of Atherosclerosis. Atherosclerosis 209, 226–229, doi:10.1016/j.atherosclerosis.2009.08.037 (2010).

16 Lakoski, S. G. et al. The relationship between blood pressure and C-reactive protein in the Multi-Ethnic Study of Atherosclerosis (MESA). Journal of the American college of cardiology 46, 1869–1874 (2005).

17 Lakoski, S. et al. The relationship between inflammation, obesity and risk for hypertension in the Multi-Ethnic Study of Atherosclerosis (MESA). Journal of human hypertension 25, 73–79 (2011).

18 Silverman, M. G. et al. Impact of Race, Ethnicity, and Multimodality Biomarkers on the Incidence of New-Onset Heart Failure With Preserved Ejection Fraction (from the Multi-Ethnic Study of Atherosclerosis). American Journal of Cardiology 117, 1474–1481, doi:10.1016/j.amjcard.2016.02.017 (2016).

19 Steffen, B. T. et al. Ethnicity, plasma phospholipid fatty acid composition and inflammatory/endothelial activation biomarkers in the Multi-Ethnic Study of Atherosclerosis (MESA). European Journal of Clinical Nutrition 66, 600–605, doi:10.1038/ejcn.2011.215 (2012).

20 Veeranna, V. et al. Association of novel biomarkers with future cardiovascular events is influenced by ethnicity: Results from a multi-ethnic cohort. International Journal of Cardiology 166, 487–493, doi:10.1016/j.ijcard.2011.11.034 (2013).

21 Weiner, S. D. et al. Systemic inflammation and brachial artery endothelial function in the Multi-Ethnic Study of Atherosclerosis (MESA). Heart 100, 862-866, doi:10.1136/heartjnl-2013-304893 (2014).

22 Wener, M. H., Daum, P. R. & McQuillan, G. M. The influence of age, sex, and race on the upper reference limit of serum C-reactive protein concentration. The Journal of rheumatology 27, 2351–2359 (2000).

23 Kottgen, A. et al. Serum cystatin C in the united states: The third national health and nutrition examination survey (NHANES III). American Journal of Kidney Diseases 51, 385–394 (2008).

24 Pan, Y. & Jackson, R. T. Ethnic difference in the relationship between acute inflammation and serum ferritin in US adult males. Epidemiology and infection 136, 421–431 (2008).

25 Berrigan, D. et al. Race/ethnic variation in serum levels of IGF-I and IGFBP-3 in US adults. Growth Hormone & IGF Research 19, 146–155 (2009).

26 Merkin, S. S. et al. Neighborhoods and cumulative biological risk profiles by race/ethnicity in a national sample of US adults: NHANES III. Annals of epidemiology 19, 194–201 (2009).

27 Juraschek, S. P. et al. Comparison of serum concentrations of β-trace protein, β2-microglobulin, cystatin C, and creatinine in the US population. Clinical Journal of the American Society of Nephrology 8, 584–592 (2013).

28 Hulver, M. W., Saleh, O., MacDonald, K. G., Pories, W. J. & Barakat, H. A. Ethnic differences in adiponectin levels. Metabolism 53, 1–3 (2004).

29 Faupel-Badger, J. M., Berrigan, D., Ballard-Barbash, R. & Potischman, N. Anthropometric correlates of insulin-like growth factor 1 (IGF-1) and IGF binding protein-3 (IGFBP-3) levels by race/ethnicity and gender. Annals of epidemiology 19, 841–849 (2009).

30 Lewontin, R. C. Ken-Ichi Kojima September 17, 1930-November 14, 1971. Genetics 71, Suppl 2:s89–90 (1972).

31 Wilson, J. F. et al. Population genetic structure of variable drug response. Nat Genet 29, 265–269, doi:10.1038/ng761 (2001).

32 Royal, C. D. & Dunston, G. M. Changing the paradigm from’race’to human genome variation. Nature genetics 36, S5–S7 (2004).

33 Owens, K. & King, M. C. Genomic views of human history. Science 286, 451–453 (1999).

34 Sidoli, S., Kulej, K. & Garcia, B. A. Why proteomics is not the new genomics and the future of mass spectrometry in cell biology. J Cell Biol 216, 21–24, doi:10.1083/jcb.201612010 (2017).

35 Schwanhausser, B. et al. Global quantification of mammalian gene expression control. Nature 473, 337–342, doi:10.1038/nature10098 (2011).

36 Berasain, C. & Avila, M. A. The EGFR signalling system in the liver: from hepatoprotection to hepatocarcinogenesis. J Gastroenterol 49, 9–23, doi:10.1007/s00535-013-0907-x (2014).

37 Soh, J. et al. Ethnicity affects EGFR and KRAS gene alterations of lung adenocarcinoma. Oncol Lett 10, 1775–1782, doi:10.3892/ol.2015.3414 (2015).

38 Phua, L. C. etal Prevalence of KRAS, BRAF, PI3K and EGFR mutations among Asian patients with metastatic colorectal cancer. Oncol Lett 10, 2519–2526, doi:10.3892/ol.2015.3560 (2015).

39 Paez, J. G. et al. EGFR mutations in lung cancer: correlation with clinical response to gefitinib therapy. Science 304, 1497–1500, doi:10.1126/science.1099314 (2004).

40 Markides, K. S. & Eschbach, K. Aging, migration, and mortality: current status of research on the Hispanic paradox. J GerontolB Psychol Sci Soc Sci 60 Spec No 2, 68–75 (2005).

41 Royall, D. R., Al-Rubaye, S., Bishnoi, R. & Palmer, R. F. Serum proteins mediate depression’s association with dementia. Plos One 12, doi:10.1371/journal.pone.0175790 (2017).

42 Royall, D. R., Palmer, R. F., O’Bryant, S. E. & Consortium, T. A. s. R. a. C. Validation of a latent variable representing the dementing process. J Alzheimers Dis 30, 639–649, doi:10.3233/JAD-2012-120055 (2012).

43 John, S. E. et al. The Effectiveness and Unique Contribution of Neuropsychological Tests and the delta Latent Phenotype in the Differential Diagnosis of Dementia in the Uniform Data Set. Neuropsychology 30, 946–960, doi:10.1037/neu0000315 (2016).

44 Gavett, B. E. et al. The delta Latent Dementia Phenotype in the Uniform Data Set: Cross-Validation and Extension. Neuropsychology 29, 344–352, doi:10.1037/neu0000128 (2015).

45 Palmer, R. F. & Royall, D. R. Future Dementia Severity is Almost Entirely Explained by the Latent Variable delta’s Intercept and Slope. Journal of Alzheimers Disease 49, 521–529, doi:10.3233/jad-150254 (2016).

46 Royall, D. R. & Palmer, R. F. δ scores predict mild cognitive impairment and Alzheimer’s disease conversions from nondemented states. Alzheimers Dement (Amst) 6, 214–221, doi:10.1016/j.dadm.2017.02.002 (2017).

47 Yu, Z. et al. Human serum metabolic profiles are age dependent. Aging Cell 11, 960–967, doi:10.1111/j.1474-9726.2012.00865.x (2012).

48 Waring, S. et al. The Texas Alzheimer’s Research Consortium longitudinal research cohort: study design and baseline characteristics. Texas Public Health Journal, 10–13 (2008).

49 O’bryant, S. E. et al. A serum protein-based algorithm for the detection of Alzheimer disease. Archives of neurology 67, 1077–1081 (2010).

50 Webb-Robertson, B. J. et al. Review, evaluation, and discussion of the challenges of missing value imputation for mass spectrometry-based label-free global proteomics. J Proteome Res 14, 1993–2001, doi:10.1021/pr501138h (2015).

51 Breiman, L. Random forests. Machine learning, 5–32 (2001).

52 Lawrence, R. L., Wood, S. D. & Sheley, R. L. Mapping invasive plants using hyperspectral imagery and Breiman Cutler classifications (RandomForest). Remote Sensing of Environment 100, 356–362 (2006).

53 Strobl, C., Boulesteix, A.-L., Zeileis, A. & Hothorn, T. Bias in random forest variable importance measures: Illustrations, sources and a solution. BMC bioinformatics 8, 25 (2007).

54 Diaz-Uriarte, R. GeneSrF and varSelRF: a web-based tool and R package for gene selection and classification using random forest. BMC Bioinformatics 8, 328, doi:10.1186/1471-2105-8-328 (2007).

55 Breiman, L., Friedman, J., Stone, C. J. & Olshen, R. A. Classification and Regression Trees. (CRC Press, 1984).

56 Prinzie, A. & Van den Poel, D. Random forests for multiclass classification: Random multinomial logit. Expert systems with Applications 34, 1721–1732 (2008).

57 Harrell, F. E., Lee, K. L. & Mark, D. B. Tutorial in biostatistics multivariable prognostic models: issues in developing models, evaluating assumptions and adequacy, and measuring and reducing errors. Statistics in medicine 15, 361–387 (1996).

58 Fawcett, T. ROC graphs: notes and practical considerations for data mining researchers Technical Report HPL-2003-4. HP Labs (2003).

59 Franceschini, A. et al. STRING v9.1: protein-protein interaction networks, with increased coverage and integration. Nucleic Acids Research 41, D808–D815, doi:10.1093/nar/gks1094 (2013).

